# Lifespan trajectories and relationships to memory of the macro- and microstructure of the anterior and posterior hippocampus – a longitudinal multi-modal imaging study

**DOI:** 10.1101/564732

**Authors:** Espen Langnes, Markus H. Sneve, Donatas Sederevicius, Inge K. Amlien, Kristine B Walhovd, Anders M Fjell

## Abstract

There is evidence for a hippocampal long axis anterior-posterior (AP) differentiation in memory processing, which may have implications for the changes in episodic memory performance typically seen across development and aging. The hippocampal formation shows substantial structural changes with age, but the lifespan trajectories of hippocampal sub-regions along the AP axis are not established. The aim of the present study was to test whether the micro- and macro-structural age-trajectories of the anterior (aHC) and posterior (pHC) hippocampus are different. In a single-center longitudinal study, 1790 cognitively healthy participants, 4.1-93.4 years of age, underwent a total of 3367 MRI examinations and 3033 memory tests sessions over 1-6 time points, spanning an interval up to 11.1 years. T1-weighted scans were used to estimate the volume of aHC and pHC, and diffusion tensor imaging to measure mean diffusion (MD) within each region. We found that the macro- and microstructural lifespan-trajectories of aHC and pHC were clearly distinguishable, with partly common and partly unique variance shared with age. aHC showed a protracted period of microstructural development, while pHC microstructural development as indexed by MD was more or less completed in early childhood. In contrast, pHC showed larger unique aging-related changes. A similar aHC – pHC difference was observed for volume, although not as evident as for microstructure. All sub-regions showed age-dependent relationships to episodic memory function. For aHC micro- and macrostructure, the relationships to verbal memory performance varied significantly with age, being stronger among the older participants. Future research should disentangle the relationship between these structural properties and different memory processes – encoding vs. retrieval in particular – across the lifespan.

## Introduction

Hippocampal sub-regions support different episodic memory processes (Collin, Milivojevic, & Doeller, 2015; Moscovitch et al., 2016; Strange, Witter, Lein, & Moser, 2014). There is convincing evidence for a long axis anterior-posterior (AP) specialization (Chase et al., 2015; Kühn & Gallinat, 2014; Poppenk, Evensmoen, Moscovitch, & Nadel, 2013), with encoding being supported relatively more by the anterior hippocampus (aHC) and retrieval relatively more by the posterior (pHC) (Kühn & Gallinat, 2014; Lepage, Habib, & Tulving, 1998; Nadel, Hoscheidt, & Ryan, 2012; Poppenk et al., 2013). The hippocampal formation shows substantial structural changes through life (Allen, Bruss, Brown, & Damasio, 2005; Walhovd et al., 2005), which in turn contributes to the lifespan trajectory of episodic memory (Fjell et al., 2014; Raz, Rodrigue, Head, Kennedy, & Acker, 2004). However, the trajectories of the hippocampal AP sub-regions in development and aging are not established. Studies assessing age effects on hippocampal volume along the AP-axis have provided evidence that different segments show different developmental (DeMaster, Pathman, Lee, & Ghetti, 2014; Lin et al., 2013; Riggins et al., 2018) and aging (Apostolova et al., 2012; Csernansky et al., 2000; Hackert et al., 2002; Wang et al., 2003; Yang, Goh, Chen, & Qiu, 2013) trajectories. Still, while it seems clear that both global (Fjell et al., 2010; Fjell, Westlye, et al., 2013) and regional (Daugherty, Bender, Raz, & Ofen, 2016) hippocampal changes are highly non-linear over age, the shape of these trajectories across the AP axis, and which AP regions that change more or less during different ages, are not known. For instance, a combination of U-shaped and inverse U-shaped age-relationships have been reported for different segments of the AP axis in development (Daugherty, Flinn, & Ofen, 2017; Gogtay et al., 2006; Schlichting, Guarino, Schapiro, Turk-Browne, & Preston, 2017). Longitudinal examinations may help resolve this issue. Further, mapping trajectories over wider age-ranges will provide higher accuracy in determining the precise d evelopmental and aging trajectory of each sub-region.

Hippocampal volume changes in development and aging could ultimately reflect various cellular processes happening within the hippocampus, such as neurogenesis (Goncalves, Schafer, & Gage, 2016) and non-neuronal cell changes (Bechmann & Nitsch, 2000), cell death and synaptic changes (Lester, Moffat, Wiener, Barnes, & Wolbers, 2017; Small, Schobel, Buxton, Witter, & Barnes, 2011), pruning (Kantor & Kolodkin, 2003), myelination (Nickel & Gu, 2018) and vascularization (Tatu & Vuillier, 2014). Several of these will likely affect water diffusion in hippocampal tissue, which can be measured by diffusion tensor imaging (DTI). Evidence so far suggests that higher mean diffusivity (MD) in the hippocampus is positively related to aging (den Heijer et al., 2012; Pereira et al., 2014; Wolf, Fischer, de Flores, Chetelat, & Fellgiebel, 2015), increases over time in older adults (Anblagan et al., 2018) and correlates more strongly with poorer memory function than hippocampal volume (Aribisala et al., 2014; Carlesimo, Cherubini, Caltagirone, & Spalletta, 2010; den Heijer et al., 2012; van Norden et al., 2012). Evidence also suggests that hippocampal MD and volume could be inversely related, with studies reporting nominal negative correlations that usually are not statistically significant (Anblagan et al., 2018; den Heijer et al., 2012; Pereira et al., 2014). Although few studies have directly tested micro- vs. macro-structural measures of hippocampal subfields, the tendency seems to be that they are related but complimentary in their aging-relationships (Pereira et al., 2014; Wolf et al., 2015). We could not find any published studies of hippocampal MD changes along the AP axis in development. Mapping microstructural changes in hippocampal AP sub-regions across development and aging, and relate these to macro-structural, volumetric changes, will yield new understanding of hippocampal changes through life.

The main aim of the present longitudinal study was to test whether the micro-and macro-structural changes of aHC and pHC followed different trajectories through life. MD and volume were quantified in 3367 MR scans (3261 for MD) from a single-center sample of 1790 (1657 for MD) cognitively healthy participants from 4.1 to 93.4 years, to address three questions:

1. Do anterior and posterior hippocampal macro- and micro-structure follow different lifespan trajectories? The existing evidence for volume is inconsistent, while micro-structure has not been mapped through lifespan.
2. What is the relationship between micro- and macro-structural hippocampal age-changes? We hypothesized that they would show mainly independent age-relationships.
3. How do aHC and pHC changes relate to episodic memory performance? Volumetric studies have yielded mixed results (Van Petten, 2004), but there are indications in favor of longitudinal compared to cross-sectional relationships (Fjell, McEvoy, et al., 2013; Gorbach et al., 2017; Tamnes et al., 2014), and that correlations may be stronger in certain hippocampal regions (Carr et al., 2017; Daugherty et al., 2017; DeMaster et al., 2014; Hackert et al., 2002; Nordin, Herlitz, Larsson, & Soderlund, 2017; Valdes Hernandez et al., 2017), reflecting specific parts of the memory process (Nadel et al., 2012; Poppenk et al., 2013; Poppenk & Moscovitch, 2011). In addition, relationships may be different in development vs. adulthood and aging (DeMaster et al., 2014). Based on the scarce evidence that exists, we hypothesized that MD would be more strongly related to memory performance than volume and that relationships may be stronger in development and aging than young adulthood and mid-life.

## Materials and methods

### Sample

Participants were drawn from studies coordinated by the Research Group for Lifespan Changes in Brain and Cognition (LCBC www.oslobrains.no) (Fjell et al., 2015), approved by a Norwegian Regional Committee for Medical and Health Research Ethics. Written informed consent was obtained from all participants older than 12 years of age and from a parent/guardian of volunteers under 16 years of age. Oral informed consent was obtained from participants under 12 years of age. The full sample consisted of 1790 healthy participants, 4.1-93.4 years of age (mean examination age, 36.2 years, 1^st^ quartile = 12.5, 3^rd^ quartile = 63.2) with a total of 3367 MRI examinations (3261 DTI) and 3033 memory tests sessions (3033 with verbal memory, 2532 with visual memory, see below). Participants were followed for up to six time points with MRI, for a maximum period of 11.13 years since baseline (mean 2.7 years, 1^st^ quartile = 0.6, 3^rd^ quartile = 3.8). Mean interval between each follow up was 2.0 years (1^st^ quartile = 0.2, 3^rd^ quartile = 3.4). Adult participants were screened using a standardized health interview prior to inclusion in the study. Participants with a history of self- or parent-reported neurological or psychiatric conditions, including clinically significant stroke, serious head injury, untreated hypertension, diabetes, and use of psychoactive drugs within the last two years, were excluded. Further, participants reporting worries concerning their cognitive status, including memory function, were excluded. All participants 40-80 years of age were required to score >26 and participants > 80 years > 25 on the Mini Mental State Examination (Folstein, Folstein, & McHugh, 1975) according to population norms (Crum, Anthony, Bassett, & Folstein, 1993). See Table 1 for sample descriptives.

**Table 1.** Sample descriptives. MMS at baseline: Available for 868. FSIQ

|  |  |
| --- | --- |
| Number of participants | 1790 |
| Number of examinations in total | 3367 |
| Hippocampal volume examinations | 3367 |
| Hippocampal volume examinations | 3261 |
| Verbal memory examinations total | 3033 |
| Verbal memory examinations with MRI | 2851 |
| Visual memory examinations total | 2532 |
| Visual memory examinations with MRI | 2417 |
| Mean number of examinations | 1.97 (1-6) |
| Mean interval between examinations | 2.7 (0.1-11.1) |
| Age at baseline | 32.0 (23.6) |
| Sex | 1056F/ 733M |
| MMS at baseline | 29.0 (SD = 1.1) |
| FSIQ | 112.0 (SD = 11.6) |

**Table 2.** Cross-scanner comparison

|  | Avanto | Skyra | % Diff | t | p= | r | p= |
| --- | --- | --- | --- | --- | --- | --- | --- |
| Anterior MD | 1.08E<sup>-03</sup> | 1.06E<sup>-03</sup> | 2.62 | 7.94 | 2.1E<sup>-13</sup> | .71 | 1.0E<sup>-28</sup> |
| Posterior MD | 1.11E<sup>-03</sup> | 1.12E<sup>-03</sup> | -0.90 | -2.12 | .036 | .73 | 4.6E<sup>-31</sup> |
| Anterior volume | 1988 | 2090 | -5.00 | -9.98 | 3.6E<sup>-16</sup> | .85 | 4.0E<sup>-52</sup> |
| Posterior volume | 2010 | 2063 | -2.60 | -5.51 | 1.2E<sup>-7</sup> | .88 | 1.3E<sup>-58</sup> |

### Memory testing

The California Verbal learning Test (CVLT) (Delis, Kramer, Kaplan, & Ober, 2000) was used to assess verbal episodic memory and the Rey-Osterrieth Complex Figure (CFT) test (Meyers & Meyers, 1995) was used to assess visual episodic memory. To allow comparison between verbal and visual memory, scores on the 30 minutes recall conditions were used for both tests.

The learning part of CVLT consists of oral presentation of 16 words in four semantically related categories, and the whole list is presented five times with a free recall trial after each presentation. After five presentations, the free recall trials are repeated 5 and 30 minutes later. To avoid fatigue in the children below 6.5 years, only three of the categories were presented. However, in the age-category 6.19-6.90 years, some participants were given the four- and some the three-category test, allowing us directly to compare the test scores. The scores were indistinguishable between test versions, and the age-slopes were as expected for both versions. Thus, the CVLT scores for both versions were entered into the statistical analyses (see below). For a subsample of the children and adolescents, Hopkins Verbal Learning Test was used. This test is based on the same sructure as the CVLT, with 4-items categories of concrete nouns to be learned by repeated presentations. HVLT consists of 3 categories (12 items in total) and 3 repetitions. Previous studies have shown good construct validity for HVLT (Shapiro, Benedict, Schretlen, & Brandt, 1999; Woods et al., 2005), and that the main measures extracted from the HVLT, such as used in the present study, show good correspondence with similar measures from the CVLT (Lacritz & Cullum, 1998; Lacritz, Cullum, Weiner, & Rosenberg, 2001). These scores were then transformed into CVLT equivalent scores by multiplying the test scores with 4/3. Validation analyses were run, confirming that the different test versions did not affect the results (see Results section below).

For visual memory, CFT measures visuo-constructive memory using a novel, complex design which participants are asked to copy and then reproduce from memory after 30 min. The participants were presented with a picture of a geometrical figure on an A4 sheet of paper and were asked to draw the figure as similar as possible. After approximately 30 min, during which time the participants completed other tasks with mainly verbal material, they were asked to draw the figure again without the original picture in front of them. The scoring system divides the figure into 18 subunits and awards 2 points for each correct and correctly placed unit, 1 point for an inaccurately drawn or incorrectly placed unit, and a 1/2 point for a unit that is recognizable but both inaccurate and inaccurately placed in the drawing. This results in a maximum score of 36 points for each drawing.

### MRI acquisition and cross-scanner validation

Imaging data were acquired on two different MRIs systems: 720 participants were scanned on a 1.5 Tesla Avanto (12 channel head coil) and 1070 on a 3T Skyra (32 channel had coil). Importantly, all longitudinal observations were from the same scanner for each participant.

Avanto T1-weighted: 2 repeated 3D T1-weighted magnetization prepared rapid gradient echo (MPRAGE): TR/TE/TI = 2400 ms/ 3.61 ms/ 1000 ms, FA = 8°, acquisition matrix 192 × 192, FOV = 240×240 mm, 160 sagittal slices with voxel sizes 1.25 × 1.25 × 1.2 mm. For most children 4-9 years old, iPAT was used, acquiring multiple T1 scans within a short scan time, enabling us to discard scans with residual movement and average the scans with sufficient quality.

Avanto DTI: 32 directions, TR = 8200 ms, TE = 81 ms, b-value = 700 s/mm^2,^ voxel size = 2.0 × 2.0 × 2.0 mm, field of view = 128, matrix size = 128 × 128 × 64, number of b0 images = 5, GRAPPA acceleration factor = 2.

Skyra T1-weighted: 176 sagittal oriented slices was obtained using a turbo field echo pulse sequence (TR = 2300 ms, TE = 2.98 ms, flip angle = 8°, voxel size = 1 × 1 × 1 mm, FOV = 256 × 256 mm). For the youngest children, integrated parallel acquisition techniques (iPAT) was used, acquiring multiple T1 scans within a short scan time, enabling us to discard scans with residual movement and average the scans with sufficient quality.

Skyra DTI: A single-shot twice-refocused spin-echo echo planar imaging (EPI) with 64 directions: TR = 9300 ms, TE = 87 ms, b-value = 1000 s/mm^2^, voxel size = 2.0 × 2.0 × 2.0 mm, slice spacing = 2.6 mm, FOV = 256, matrix size = 128 × 130 × 70, 1 non-diffusion-weighted (b = 0) image. A bB0-weighted image was acquired with the reverse phase encoding.

Since different scanners will yield different volumetric and MD values, 180 participants evenly distributed across the age range were scanned on the 1.5T (Avanto) scanner and the 3T (Skyra) scanner on the same day, to allow us to directly assess the influence of scanner. As expected, the different scanners yielded highly significant differences in absolute MD and volume (Table 3). The correlations between scanners were good, however, r = .85 (anterior) and .88 (posterior) for volume and .71 (anterior) and .73 (posterior) for MD. The high rank-order coherence between scanners suggested that inclusion of scanner as a covariate in the analyses efficiently would remove most of the variance between participants due to different scanners. In addition, validation analyses were run, replicating the main findings with data from each scanner separately (see Results section below).

**Table 3.** Relationship to age. GAMM with all brain variables as joint predictors of age was run, with sex, scanner and ICV as covariates of no interest. Edf: effective degrees of freedom. MD: Mean diffusion.

|  | F | edf | p ≤ |
| --- | --- | --- | --- |
| Anterior volume | 24.2 | 1.0 | 9.18e<sup>-07</sup> |
| Posterior volume | 41.8 | 5.0 | 2e<sup>-16</sup> |
| Anterior MD | 2.9 | 4.5 | 0.01 |
| Posterior MD | 37.6 | 4.6 | 2e<sup>-16</sup> |

### MRI preprocessing - morphometry

T1-weighted scans were run through FreeSurfer 6.0 (https://surfer.nmr.mgh.harvard.edu/). FreeSurfer is an almost fully automated processing tool (Dale, Fischl, & Sereno, 1999; Fischl et al., 2002; Fischl, Sereno, & Dale, 1999; Fischl, Sereno, Tootell, & Dale, 1999), and manual editing was not performed to avoid introducing errors. For the children, the issue of movement is especially important, as it could potentially induce bias in the analyses (Reuter et al., 2015). Rather, all scans were manually rated for movement on a 1-4 scale, and only scans with ratings 1 and 2 (no visible or only very minor possible signs of movement) were included in the analyses, reducing the risk of movement affecting the results. Also, all reconstructed surfaces were inspected, and discarded if they did not pass internal quality control. The hippocampus was initially segmented as part of the FreeSurfer subcortical stream (Fischl et al., 2002) before being divided in aHC and pHC (see below).

### MRI preprocessing – DTI

DTI scans were processed with FMRIB’s Diffusion Toolbox (fsl.fmrib.ox.ac.uk/fsl/fslwiki) (Jenkinson, Beckmann, Behrens, Woolrich, & Smith, 2012; Smith et al., 2004). B0 images were also collected with reversed phase-encode blips, resulting in pairs of images with distortions going in opposite directions. From these pairs we estimated the susceptibility-induced off-resonance field using a method similar to what is described in (Andersson et al., 2003) as implemented in FSL (Smith et al., 2004). We then applied the estimate of the susceptibility induced off-resonance field with the eddy tool (Andersson and Sotiropoulos, 2016), which was also used to correct eddy-current induced distortions and subject head movement, align all images to the first image in the series and rotate the bvecs in accordance with the image alignments performed in the previous steps (Jenkinson et al., 2002; Leemans and Jones, 2009).

### Hippocampal anterior-posterior segmentation

Moving anteriorly through the coronal planes of an MNI-resampled human brain, y = −21 corresponds to the appearance of the uncus of the parahippocampal gyrus. In line with recent recommendations for long-axis segmentation of the hippocampus in human neuroimaging (Poppenk et al., 2013), we labeled hippocampal voxels at or anterior to this landmark as anterior HC while voxels posterior to the uncal apex were labeled as posterior HC. Specifically, for each participant, all diffusion voxels for which more than 50% of the underlying anatomical voxels were labeled as hippocampus by FreeSurfer (Fischl et al., 2002) were considered representations of the hippocampus. While keeping the data in native subject space, we next established hippocampal voxels’ locations relative to MNI y = −21 by calculating the inverse of the MNI-transformation parameters for a given subject’s brain and projecting the back-transformed coronal plane corresponding to MNI y = −21 to diffusion native space. All reported diffusion measures thus represent averages from hippocampal sub-regions established in native space. An illustration of this segmentation is shown in Figure 1.

**Figure 1.**
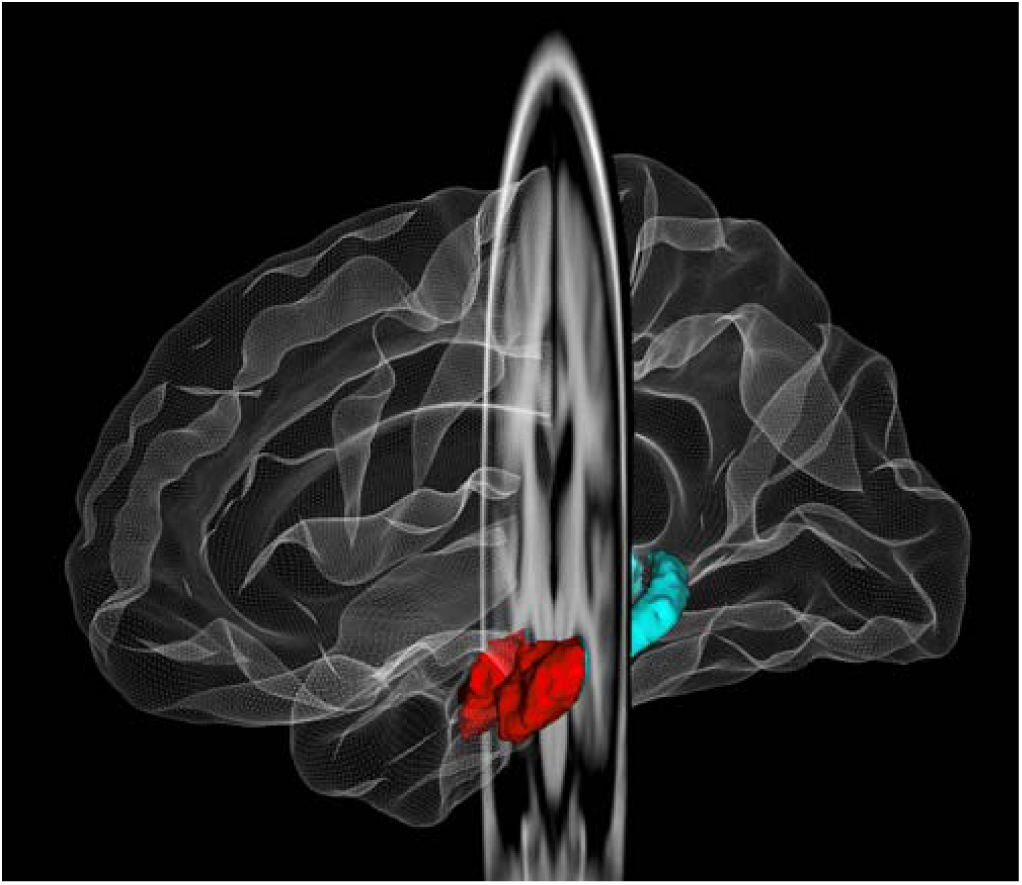
Segmentation of the hippocampus

### Statistics

Analyses were run in R (https://www.r-project.org) using Rstudio (www.rstudio.com) IDE. Generalized Additive Mixed Models (GAMM) using the package “mgcv” (Wood, 2006) were used to derive age-functions. Each hippocampal sub-region was used as dependent variable in separate analyses. We included a smooth term for age, random effect for subject, and sex and scanner as covariates of no interest in all analyses of hippocampal sub-regions. For volume, estimated intracranial volume (ICV) was also included in the model. To test whether different sub-regions (aHC, pHC) and metrics (volume, MD) were uniquely related to age, and to derive residualized age-functions of each measure independently of the other measures, GAMMs were run with age as dependent variable and each of the imaging measures of interest as joint predictors, with the same covariates as above. To test the relationshi pbetween the hippocampal variables and memory, models were first run with memory as dependent and age as predictor. Age was then replaced with each hippocampal variable in separate analyses. To test the hypothesized age-dependent hippocampus-memory relationship, models were run including an age × hippocampus interaction term. Subject timepoint was included as covariate in all memory analyses to control for practice effects of the memory scores.

Akaike Information Criterion (AIC) (Akaike, 1974) and the Bayesian Information Criterion (BIC) was used to guide model selection and help guard against over-fitting. The smoothness of the age-curve is estimated as part of the model fit, and the resulting effective degrees of freedom (edf) was taken as a measure of deviation from linearity. The p-values associated with the smooth terms are only approximate, as they are based on the assumption that a penalized fit is equal to an unpenalized fit with the same edf, and do not take into account uncertainty associated with the smoothing parameter estimation. The major advantage of GAMM in the present setting is that relationships of any degree of complexity can be modelled without specification of the basic shape of the relationship, and GAMM is thus especially well-suited to map life-span trajectories of neurocognitive variables which can be assumed to be non-linear and where the basic form of the curve is not known (Fjell et al., 2010). The main analyses were repeated for each scanner separately as an additional validation. Main analyses were also run with only the ordinary CVLT scores included to confirm that the different verbal memory test versions did not bias the results.

## Results

### Do anterior and posterior hippocampal macro-and micro-structure follow different trajectories through life?

GAMM curves illustrating the age-trajectories for MD and volume for aHV and pHC are presented in Figure 2. MD was highly related to age (anterior: F =89.3, edf = 8.23, p < 2e^−16^; posterior: F = 144.4 edf = 5.63, p < 2e^−16^), showing the expected non-linear age-trajectories. Still, there were clear differences between aHC and pHC. While the posterior section did not show age-effects until about 40 years, MD in aHC was greatly reduced in the same age-period, displaying a protracted developmental phase. Both aHC and pHC showed substantial increases in the last half of the age-span, but the increase started a decade later for the anterior (≈50 years) compared to the posterior (≈40 years). Comparing model fits, the model including aHC was superior according to all tested fit measures (aHC vs pHC AIC −55978 vs. −54595, BIC −559345 vs. −54553, logLik 27996 vs. 27305). Directly contrasting their age-trajectories, we included both as smooth predictors in the same model with age as dependent variable. Both were uniquely related to age, although the relationship was stronger for pHC (F= 27.0, edf = 4.16, p < 2e^−16^) than aHC (F = .1, edf = 4.2, p = .016), and this model was significantly better than either of the two models with only one sub-region (Log ratio = 77469, p < .0001 vs. the best single sub-region model). Inspection of the residual plots (Figure 3) confirmed the impression from the first set of analyses. These clearly demonstrated that the microstructure of pHC was highly aging-sensitive compared aHC, as evidenced by the positive relationship between residualized MD and age in pHC. In contrast, aHC was more strongly related to development than pHC, as evidenced by the negative relationship between residualized MD and age in this region.

**Figure 2.**
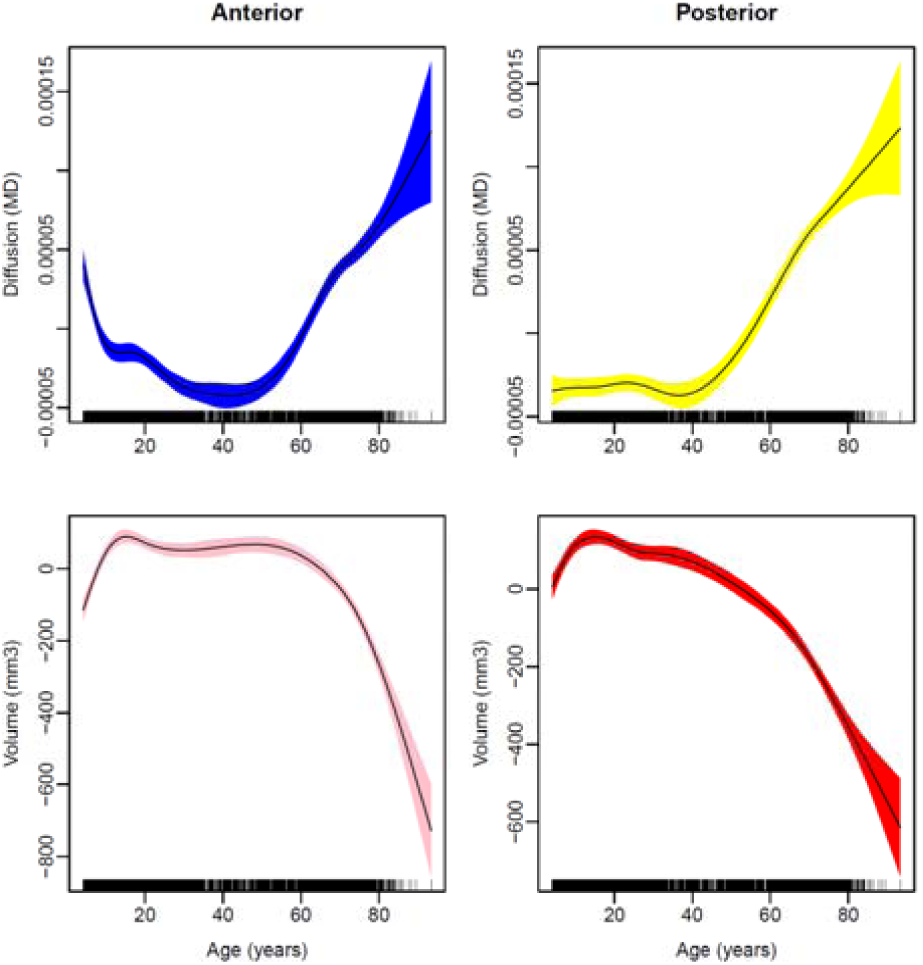
Age-trajectories for hippocampal sub-regions. The y-axis values are the residual mean diffusion/ volume where the covariates in each analysis are accounted for.

**Figure 3.**
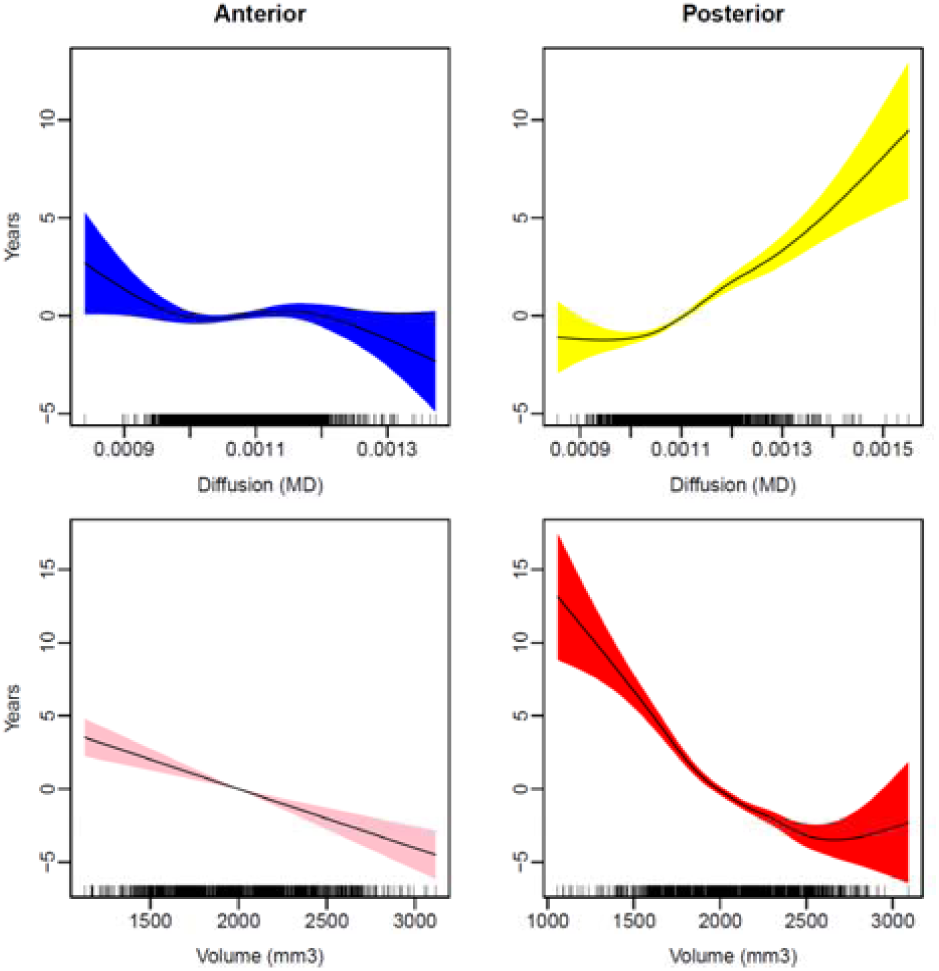
Unique age-trajectories for each sub-region. The figure depict the relationship to age (y-axis) from each sub-regions when aHC and pHC variables were included as predictors of age in the same GAMM model. The curves thus signify the unique relationship of each sub-region to age accounting for the common variance shared with the other sub-region.

The same analyses were run for hippocampal volume, with ICV as an additional covariate of no interest. As expected, volume of both sub-regions showed highly significantly inverse U-shaped age-trajectories (anterior: F = 54.2, edf = 8.11, p < 2e^−16^; posterior: F = 97.6, edf = 7.54, p < 2e^−16^). As for MD, model fit was better for the model with the anterior compared to the posterior region (anterior vs. posterior: AIC 42829 vs. 42969, BIC 42884 vs. 43024, logLik −21406 vs. −21475, respectively). Including anterior and posterior in the same model, both were still significantly related to age (anterior: F = 30.0, edf = 1.0, p = 4.57e^−08^; posterior: F = 33.5, edf = 4.99, p < 2e^−16^), and this model was significantly better than either of the two models with only one sub-region (Log ratio = 19321, p < .0001 vs. the best single sub-region model). Inspecting the residualized trajectories showed that smaller volume was linearly associated with higher age in the anterior hippocampus and non-linearly in the posterior, suggesting lower rates of residual change in the posterior hippocampus among adolescents and young adults who had the largest volumes. Having demonstrated that anterior and posterior hippocampus showed unique developmental- and aging trajectories for micro- (MD) and macro-structure (volume), we went on to test how the two modalities were interrelated in their age-trajectories.

### What is the relationship between micro-and macro-structural hippocampal changes?

To test whether macro-structural age-changes could partly be accounted for by micro-structural changes, we included both modalities in the same GAMM, with age as dependent variable and ICV as an additional covariate of no interest. For both sub-regions, MD and volume were related to age independently of each other (aHC MD: F =12.7, p = 2.45e^−11^, volume: F =79.2, p < 2e^−16^/ pHC: MD: F = 44.5, p < 2e^−16^, volume F = 46.9, p < 2e^−16^). GAMMs with each modality tested separately yielded numerically lower F-values (aHC MD F = 11.2 vs 12.7, volume F = 74.7 vs. 79.2/ pHC MD F = 34.2 vs. 44.5, volume F = 39.9 vs. 46.9 for GAMMs with separate vs. joint terms, respectively), suggesting that the age-relationships of the two modalities are independent. Thus, in a final GAMM, we included both modalities (MD, volume) and both regions (aHC, pHC) simultaneously. All were still significant predictors of age (Table 4). Plotting the residuals yielded almost identical results to those presented in Figure 3.

**Table 4.**
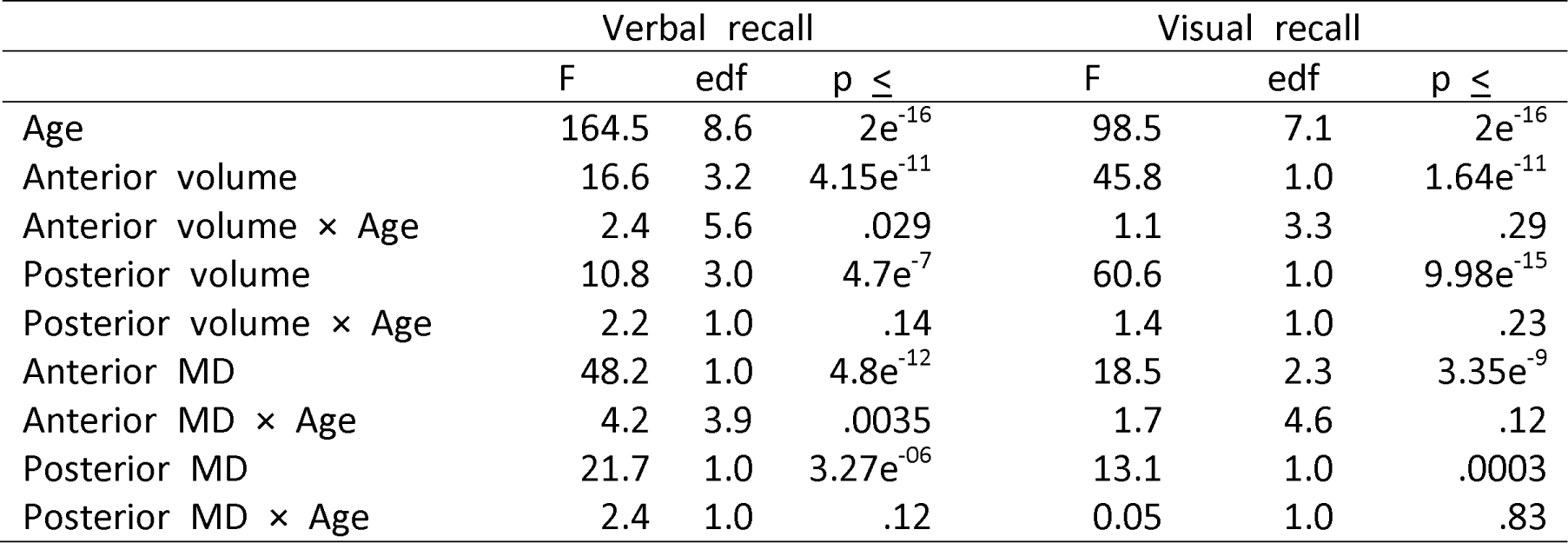
Hippocampus-memory relationships Each line represents the results of a separate GAMM. With memory as dependent variable, age and each hippocampal measure were first tested in separate GAMMs, and then additional GAMMS were run to test the age × hippocampus interaction, with age, sex, scanner, subject time-point and ICV (for volume) as covariates of no interest. Edf: effective degrees of freedom. MD: Mean diffusion. ns: not significant (p >. 05)

|  | Verbal recall |  |  | Visual recall |  |  |
| --- | --- | --- | --- | --- | --- | --- |
|  | F | edf | p ≤ | F | edf | p ≤ |
| Age | 164.5 | 8.6 | $2e^{-16}$ | 98.5 | 7.1 | $2e^{-16}$ |
| Anterior volume | 16.6 | 3.2 | $4.15e^{-11}$ | 45.8 | 1.0 | $1.64e^{-11}$ |
| Anterior volume × Age | 2.4 | 5.6 | .029 | 1.1 | 3.3 | .29 |
| Posterior volume | 10.8 | 3.0 | $4.7e^{-7}$ | 60.6 | 1.0 | $9.98e^{-15}$ |
| Posterior volume × Age | 2.2 | 1.0 | .14 | 1.4 | 1.0 | .23 |
| Anterior MD | 48.2 | 1.0 | $4.8e^{-12}$ | 18.5 | 2.3 | $3.35e^{-9}$ |
| Anterior MD × Age | 4.2 | 3.9 | .0035 | 1.7 | 4.6 | .12 |
| Posterior MD | 21.7 | 1.0 | $3.27e^{-06}$ | 13.1 | 1.0 | .0003 |
| Posterior MD × Age | 2.4 | 1.0 | .12 | 0.05 | 1.0 | .83 |

### How do aHC and pHC relate to episodic memory performance?

Both verbal and visual recall was strongly related to age, shaping inverse U-shaped relationships (Figure 4, Table 5). Separate GAMMs were run with verbal memory performance as dependent variable, and each of the brain variables as smooth predictors, with sex, scanner, subject timepoint to control or retest effects and ICV (for volume) as covariates of no interest. Hippocampal regions were strongly related to memory (all p’s <. 0001) (see Figure 5). MD was negatively related to memory score, mostly in a linear fashion, with edf = 1 for all relationships except between aHC MD and visual recall, which was characterized by an inverted U-shape. For volume, the relationships were positive - linearly for MD and inverted U-shaped for verbal recall. We re-ran the model including all sub-regional variables simultaneously. For verbal recall, aHC MD (F = 14.5, edf = 1.0, p = .00014), aHC volume (F = 14.9, edf = 2.9, p = 5.56e^−09^) and pHC volume (F = 7.9, edf = 1.0, p = .005) were still significantly related to verbal memory, while pHC MD (F = 2.5, edf = 1.0, p = .11) was not. For visual recall, significant relationships were seen for the same sub-regions, i.e. aHC MD (F = 7.5, edf = 1.0, p = .006), aHC volume (F = 6.1, edf = 1, p = 2.18e^−09^) and pHC volume (F = 31.9, edf = 1.0, p = 1.79e^−08^), while pHC MD was not significant (F = 1.5, edf = 1.0, p = .22). In contrast to verbal recall, visual recall performance seemed more strongly related to volume than MD.

**Figure 4.**
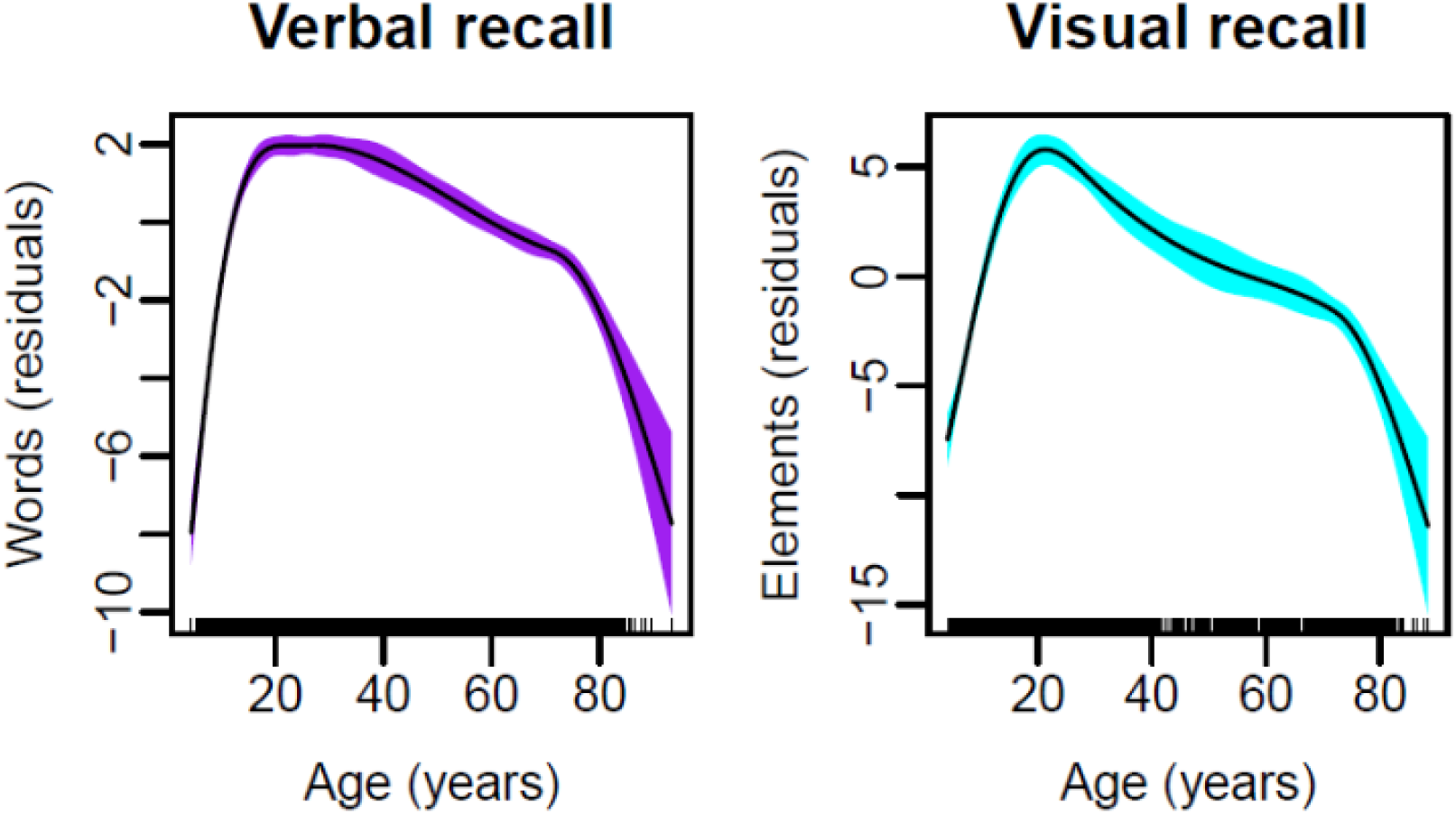
Age-trajectories of memory. Left panel: Verbal 30 min recall. Right panel: Visual 30m min recall.

**Figure 5.**
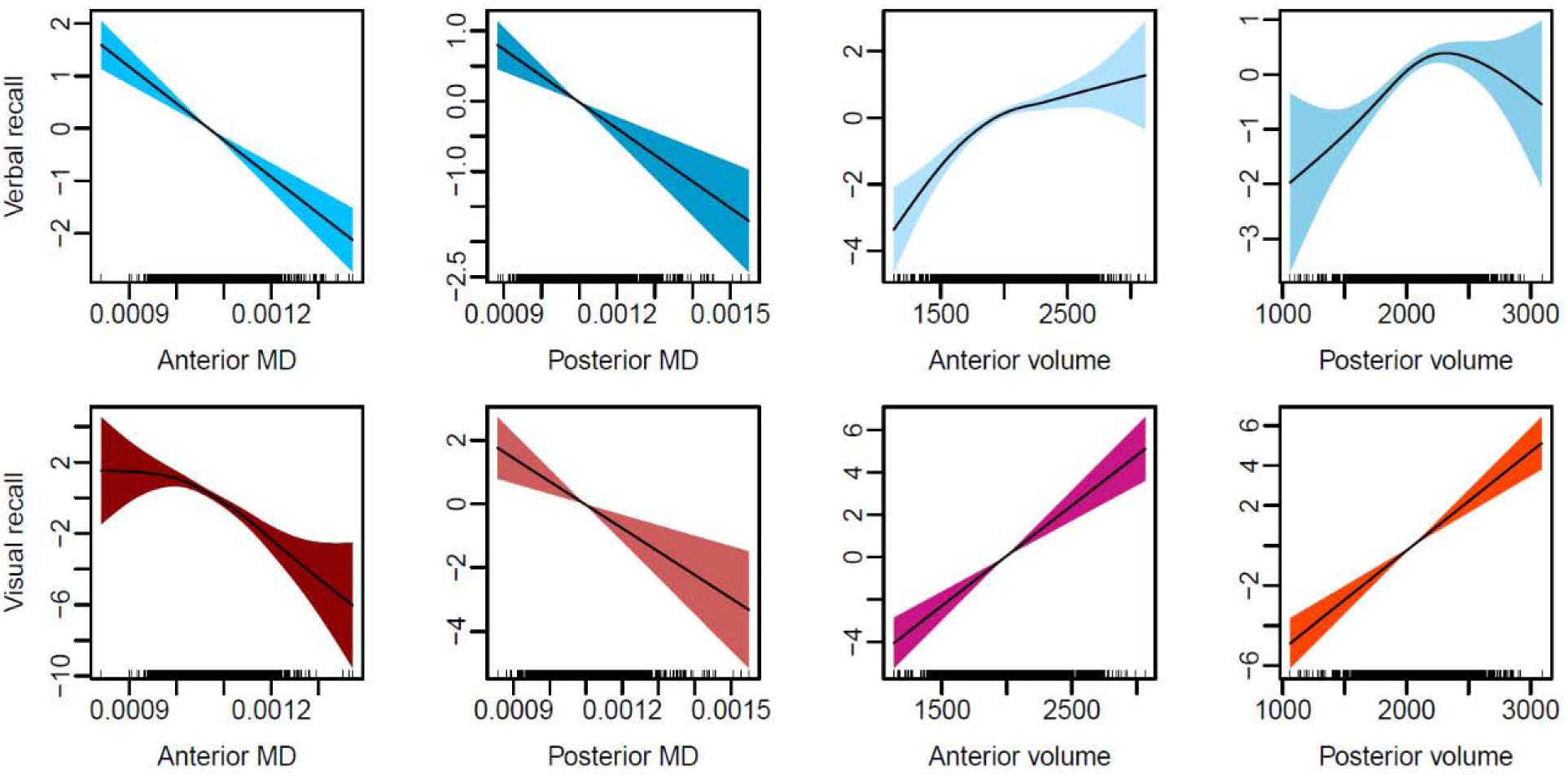
Relationships between hippocampal sub-regions and memory. Top row: Verbal memory. Bottom row: Visual memory. The y-axis values are the residual memory scores where the covariates in each analysis are accounted for.

To test the influence of age on these relationships, the analyses were repeated, first by including age as a smooth covariate, and second by including the age × hippocampus interaction term (Table 5). None of the sub-region variables predicted verbal or visual memory performance independently of age (all p’s > .052). However, for verbal recall, aHC MD and volume showed significant age-interactions. As can be seen from Figure 6, larger aHC volume was related to better memory in older adults, while smaller or no effects were seen for the adolescents, young- and middle-aged adults. For aHC MD, negligible effects were seen in middle age, with the relationships being stronger among the older participants. However, the age interactions were not particularly strong, with no p-values < .001. For visual recall, there were no significant age × hippocampus interactions.

**Figure 6.**
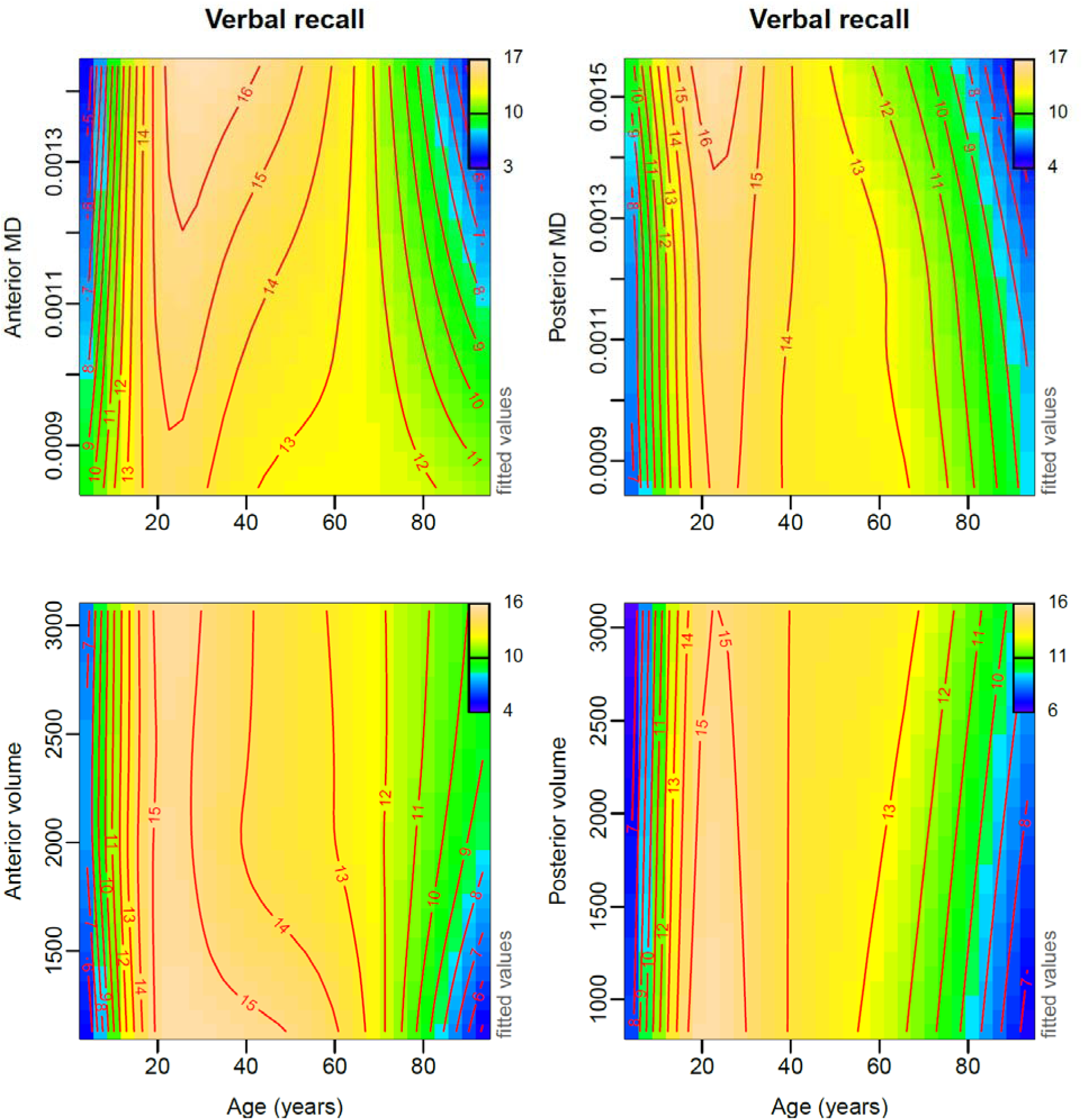
Age × hippocampus interactions for memory. There were significant age × aHC MD and age × aHC volume interactions in prediction of verbal memory (left column), but not for pHC MD or volume (right column). The colors of the contour plots signify memory score.

### Validation analyses

Although scanner was used as covariate in all analyses, we repeated the main analyses within each scanner only as an additional security measure that scanner differences did not influence the results (Avanto n = 720, 1611 T1 / 1615 DTI scans; Skyra n = 1070, 1936 T1 / 1825 DTI), see SI for details. These confirmed the age-trajectories from the full sample for each scanner separately. Joint GAMMs with MD and volume included in the same model confirmed the complementary relationships to age for both aHC and pHC for both Avanto and Skyra, except aHC MD for Skyra which only showed a trend (p = .087) (see SI for details). Finally, analyses were run testing the relationships to memory. For both scanners, the age-dependent relationships to memory performance was confirmed for all tested variables except pHC volume for the Avanto which only showed a trend (p < .056). The other 15 relationships (2 regions × 2 modalities × 2 scanners × 2 memory tests) were all significant (p < .05). Seen together with the analyses showing the high rank-order correspondence between the results from the two scanners, we believe these within-scanner analyses make a strong case that scanner differences did not influence the results.

Further, verbal memory was tested against age for only the participants with regular CVLT scores (n = 1574, 2924 examinations), excluding those with HVLT or the three-category CVLT version. The age-trajectory was close to identical to the one from the full sample (p < 2e^−16^) (see SI). The relationships between CVLT score and the sub-regions were also tested, confirming the significant age-dependent relationships (all p’s ≤ 1.26e^−05^). This shows that the current approach of jointly analyzing three different versions of the verbal recall tests did not bias the results.

## Discussion

The main discovery was that aHC and pHC showed distinguishable lifespan trajectories and age-dependent relationships to memory performance. Especially evident for MD, aHC showed a protracted developmental period of change, while pHC changed little during development. While no substantial changes in pHC MD was seen until 40 years, changes were even more evident in the last half of the age-span. This suggests that pHC microstructural development as indexed by MD from conventional DTI is more or less completed in early childhood. The same main aHC – pHC distinction was observed for macrostructure, although not to the same degree. While aHC showed a linear residual age-relationship throughout almost 9 decades of life, pHC residual changes seemed to happen mostly from middle-age and upward. Interestingly, aHC showed age-varying relationships to verbal memory function, while this was not seen for pHC. Implications of the findings are discussed below.

### Lifespan trajectories of anterior and posterior hippocampus

Both aHC and pHC volume and MD were strongly and independently related to age through the lifespan, with better model fits for aHC suggesting a somewhat tighter relationship with age. While studies of adults have shown increases in hippocampal MD with age (Anblagan et al., 2018); den Heijer et al. (2012); (Pereira et al., 2014; Wolf et al., 2015), we are not aware of any lifespan studies of hippocampal microstructure. The volumetric findings are in line with previous developmental studies showing differential (DeMaster et al., 2014; Lin et al., 2013; Riggins et al., 2018; Wang et al., 2003; Yang et al., 2013) and non-linear (Daugherty et al., 2017; Gogtay et al., 2006) age-relationships for sub-regions along the AP axis. While age-relationships have been observed for all sub-regions in some studies (Apostolova et al., 2012; Hackert et al., 2002)), others report selective or higher vulnerability of the head (Wang et al., 2003; Yang et al., 2013) or body (Malykhin, Huang, Hrybouski, & Olsen, 2017). There are also divergent findings regarding the shape of the trajectory, especially in development. Previous studies have reported both U-shaped (Daugherty et al., 2017) and a combination of U-shaped and inverse U-shaped age-relationships when moving along the AP axis (Gogtay et al., 2006; Schlichting et al., 2017). The present results clearly suggest volumetric increases in both aHC and pHC throughout childhood, forming the first segment of an inverse U-shaped lifespan trajectory. Strengths of the present study include a wide age-range covering almost 9 decades of life, a longitudinal design, decent sample size and a statistical approach capable of capturing local variations in the age-trajectories across the age-span sampled (Fjell et al., 2010; Wood, 2006). We will argue that from a life-span perspective, the general inverse U-shaped trajectory observed in the present study is in accordance with expectations.

The major objective was to disentangle the life-span trajectories of aHC and pHC micro-and macrostructure. As hypothesized, each sub-region and modality showed unique age-relationships. Interestingly, we observed a principal divide between aHC and pHC both for macro-and microstructure, most clearly manifested for the latter. While aHC showed the expected age-reduction during development and the first part of adult life, pHC did not change until mid-life (≈ 40 years). This divide was especially evident from the residual plots, where aHC and pHC showed opposite age-trajectories. In these residual analyses, age-variance common to aHC and pHC was accounted for, and the unique age-relationship of each sub-region thus isolated. In the volumetric analyses, while aHC and pHC both showed the expected inverse U-shaped age-trajectory, the joint analysis revealed an aHC vs. pHC divide with similarities to MD. The unique pHC volumetric changes were observed for higher age only, suggesting that these characterized adult and aging-related processes. aHC showed a linear negative age-relationship, thus not restricted to adult age. In contrast to microstructure, the developmental increase in volume was shared between aHC and pHC, such that neither showed unique developmental patterns.

### Independence between micro-and macro-structural lifespan changes

The inverse of the MD trajectories showed similarities to the volume trajectories. Thus, one could speculate that they would be related to some extent. As argued above, hippocampal macro-structural changes could reflect a range of cellular processes within the hippocampus potentially affecting water diffusion and thus be detectable by DTI. These could include loss of cellular barriers and increase in extracellular water content as a function of neurodegeneration (Basser & Pierpaoli, 1996; Kale et al., 2006; Pierpaoli & Basser, 1996), neurogenesis and non-neuronal cell changes, cell death and synaptic changes, pruning, myelination and vascularization (Bechmann & Nitsch, 2000; Goncalves et al., 2016; Kantor & Kolodkin, 2003; Lester et al., 2017; Nickel & Gu, 2018; Small et al., 2011; Tatu & Vuillier, 2014). Still, the results yielded no indications that hippocampal atrophy even partly could be explained by microstructural changes. The age-relationships of volume and MD were not weakened the least by including both in the same model. Even though not previously addressed in a lifespan sample, this finding is in line with previous studies showing that micro- and macrostructural measures of hippocampal subfields are complementary in their aging-relationships (Pereira et al., 2014; Wolf et al., 2015). Thus, the cellular processes responsible for the micro-and macro-structural changes in hippocampal sub-regions across life appear to be completely unrelated.

### Hippocampal structural properties and episodic memory through the lifespan

Previous studies of the hippocampus suggest that MD may be more closely associated with memory function than volume (Aribisala et al., 2014; Carlesimo et al., 2010; den Heijer et al., 2012; van Norden et al., 2012). Here, we were interested in how the memory-relationships were affected by age, and whether the contributions from aHC and pHC could be distinguished. When all sub-region variables were included in the same model, all except pHC MD contributed uniquely to explain memory performance. These results must be interpreted within a context of age changes, as the major part of the variance was shared between age, sub-region variables and memory. For aHC (MD and volume), the relationship with verbal memory performance varied as a function of age. Larger sub-region volume was related to better memory in older adults, above 60-70 years, with smaller or no effects seen in the other parts of the age-span. For MD, similar effects in the opposite direction were found. Thus, aHC showed an age-varying relationship to verbal memory while pHC did not, regardless of whether micro- or macrostructure is measured.

Importantly, however, the relative contributions from aHC vs. pHC to memory function will likely depend on the nature of the task. aHC tends to be more involved in encoding and pHC in retrieval of episodic memories (Kühn & Gallinat, 2014; Lepage et al., 1998; Nadel et al., 2012; Poppenk et al., 2013). The relative involvement of hippocampal sub-regions along the AP axis likely depends on a number of features related to the specific demands of the memory operations (Chase et al., 2015; Kühn & Gallinat, 2014; Poppenk et al., 2013). We could not distinguish between contributions from encoding vs. retrieval, which are both affected by age (Grady, 2012). In a study of hippocampal activity during encoding and retrieval of associative memories, we observed that children engaged pHC more than aHC, while aHC was more activated relative to pHC already in teenagers (Langnes et al., 2018). Thus, the differential age-trajectories of the sub-regions observed here may have implications for memory processing, which may either be disguised in the less specific memory tasks used, or detectable on the level of brain activity only. While memory-relationships with both sub-regions have been reported (Driscoll et al., 2003), studies have suggested that protracted development of hippocampal sub-regions contribute to age-related differences in episodic memory function (DeMaster et al., 2014). Similar to (DeMaster et al., 2014), we found hippocampal sub-regions to show unique relationships to memory performance that varied as a function of age, even though this was mostly evident for the oldest part of the sample in the present study. For an in-depth review and discussion of age-effects on hippocampal subfields and relationship to memory, see (de Flores, La Joie, & Chetelat, 2015).

While the age-dependent hippocampus – memory performance relationships were similar for verbal and visual recall, there were also some notable differences. First, although we did not test a formal interaction, sub-region volumes tended to be more strongly related to visual memory while MD was more strongly related to verbal memory. Second, we did not observe age-interactions for visual recall.

The reason for these differences are not clear. The visual task has a strong constructive element, related to visuo-constructive abilities (Ostby, Tamnes, Fjell, & Walhovd, 2012). The verbal task with its inherent verbal categorization is likely related to general verbal abilities. Thus, the partly different relationship to the hippocampal sub-regions and the age-interactions may result from the different basic cognitive processed involved in addition to the episodic memory component.

## Conclusion

Here we show that the micro- and macrostructure of hippocampal sub-regions along the anterior-posterior axis have distinguishable life-span trajectories, and that they are independently related to episodic verbal memory performance in an age-dependent and partly age-varying way. Future research should try to disentangle the relationship between these structural properties and different memory processes – encoding vs. retrieval in particular – across the lifespan.

## Acknowledgement

This work was supported by the Department of Psychology, University of Oslo (to K.B.W., A.M.F.), the Norwegian Research Council (to K.B.W., A.M.F.) and the project has received funding from the European Research Council’s Starting/ Consolidator Grant schemes under grant agreements 283634, 725025 (to A.M.F.) and 313440 (to K.B.W.).

