## Supplemental Information for "Lifespan trajectories and relationships to memory of the macro- and microstructure of the anterior and posterior hippocampus – a longitudinal multi-modal imaging study"

Running title (max 60 characters): Hippocampal long-axis structural changes through life

Espen Langnes^1^, Markus H. Sneve^1^, **Donatas Sederevicius^1^,** Kristine B Walhovd^1,2^, Anders M Fjell^1,2^

1) Center for Lifespan Changes in Brain and Cognition, University of Oslo, Norway

2) Department of Radiology and Nuclear Medicine, Oslo University Hospital, Oslo, Norway

Supplemental analyses separately for each scanner

**Age-trajectories for each hippocampal sub-region based on the Avanto scans only, n = 720, 1611 scans for volume/ 1615 for DTI**

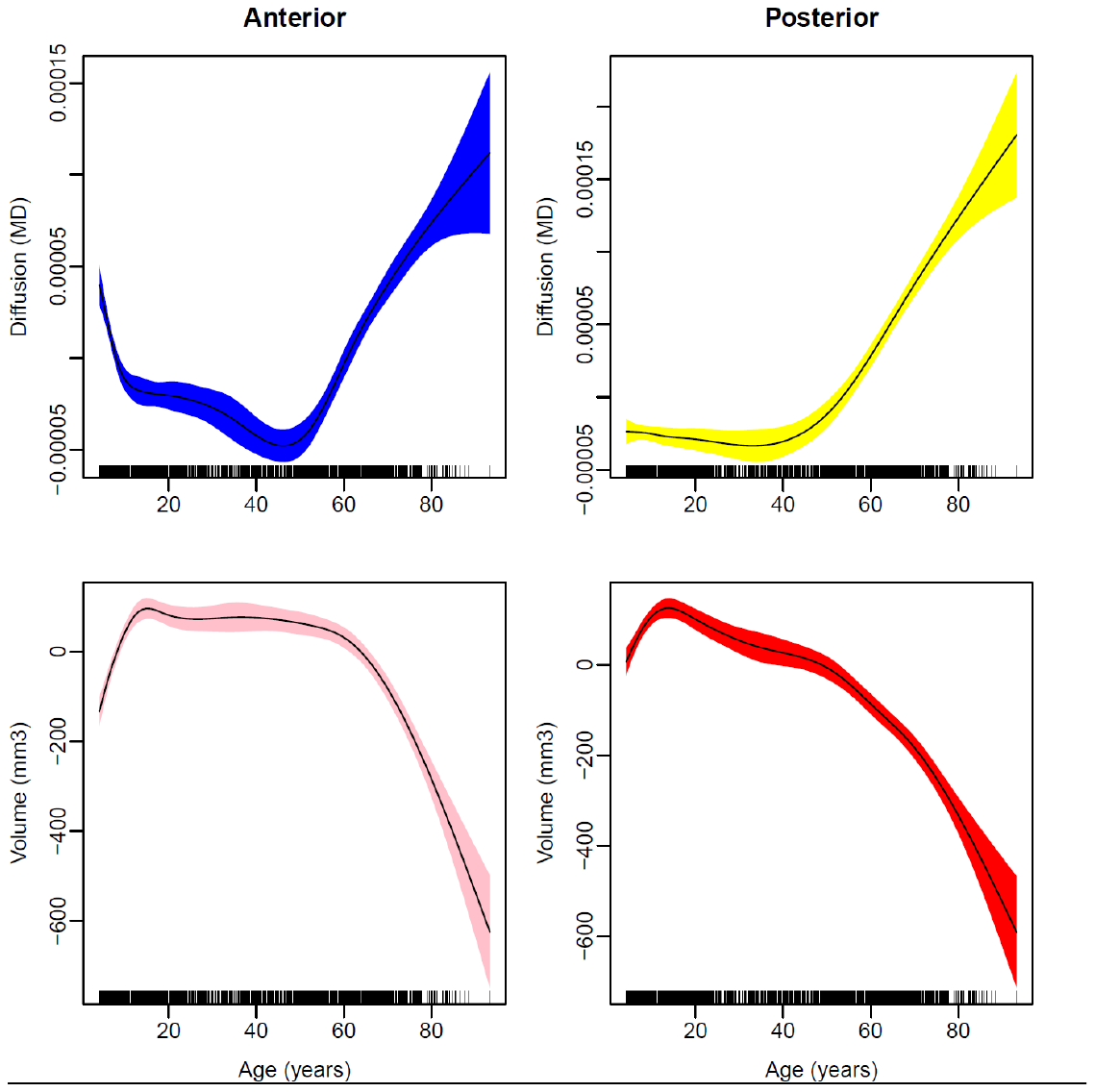

**Age-trajectories for each hippocampal sub-region based on the Skyra scans only, n = 1070, 1935 scans for volume/ 1825 for DTI**

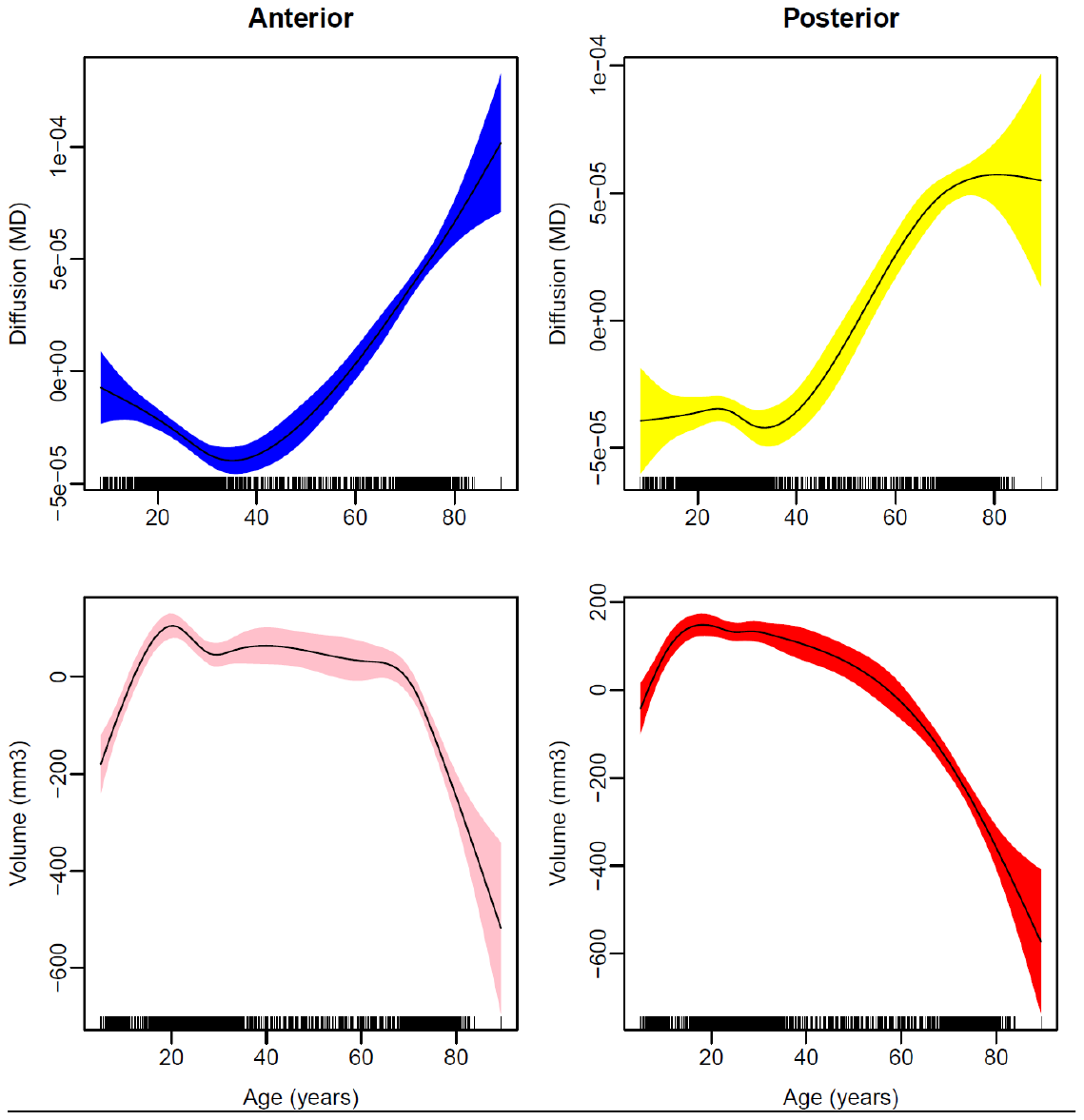

| Sub-region | MD | |  | Volume | | |
| --- | --- | --- | --- | --- | --- | --- |
|  | edf | F | p | edf | F | p |
| *Avanto* |  |  |  |  |  |  |
| aHC | 2.9 | 12.5 | 6.24e^-08^ | 1.0 | 34.5 | 5.03e^-09^ |
| pHC | 4.1 | 43.1 | <2e^-16^ | 5.3 | 34.7 | <2e^-16^ |
| *Skyra* |  |  |  |  |  |  |
| aHC | 3.7 | 2.4 | .087 | 1.0 | 54.5 | 2.33e^-13^ |
| pHC | 3.9 | 3.8 | .005 | 4.1 | 17.8 | 1.75e^-14^ |

Unique age-relationships of hippocampal sub-regions microstructure vs. macrostructure. The models were fitted with GAMMs, and sex, ICV and scanner were included as covariates.

Edf: effective degrees of freedom

**Age-trajectories for verbal memory based on CVLT only (n = 1574, 2924 memory tests)**

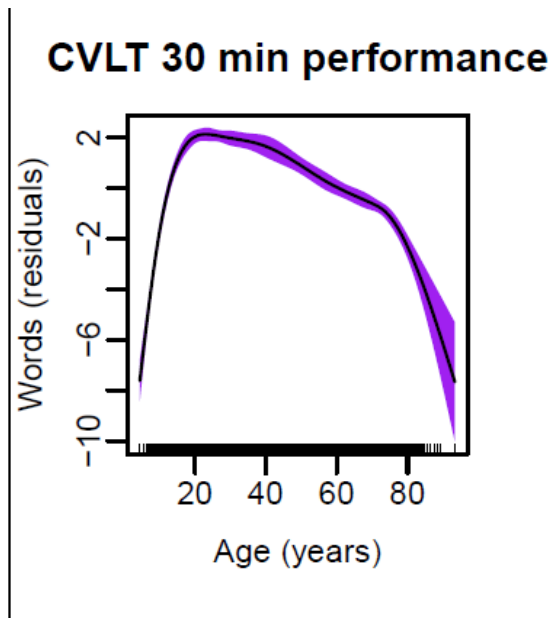
